# A streamlined and abundant *Nitrosopumilus* population retains vitamin biosynthesis under high purifying selection in the Baltic Sea

**DOI:** 10.64898/2026.10.02.756262

**Authors:** Armando Pacheco-Valenciana, Daan R. Speth, Gerrit Wienhausen, Stefan Bruns, Heinz Wilkes, Anders F. Andersson, A. Murat Eren, Sarahi L. Garcia

**Author notes:** Corresponding author: Armando Pacheco-Valenciana, Sarahi L. Garcia.

## Abstract

Genome streamlining is expected to favor the loss of costly biosynthetic functions, raising the question of what benefit justifies the continued maintenance of complex pathways in small genomes. Here, we investigated this question in an abundant *Nitrosopumilus* population from the Baltic Sea that combined a streamlined genome (1.4 Mbp) with substantial biosynthetic capacity for vitamins B_1_, B_7_, and B_12_. We found three lines of evidence that support the biological significance of this retention. First, phylogenomic analysis of 1,077 archaeal species representatives showed that B-vitamin biosynthesis is broadly conserved across ammonia-oxidizing archaea (AOA), indicating maintenance over deep evolutionary timescales. Second, genes associated with B-vitamin biosynthesis tended to be under stronger purifying selection than the genomic background across Baltic Sea metagenomes, consistent with selection preserving these functions in current environmental populations. Third, measurements in the cultured relative *Nitrosopumilus maritimus* showed B-vitamins and related precursors in both intracellular and extracellular fractions, providing a potential route through which these retained functions could influence metabolite availability beyond the producing cell. Together, our findings suggest that retained biosynthetic pathways in highly streamlined microorganisms reveal functions that are beneficial for the success of a natural population, while also pointing to their potential importance for other community members coexisting within the same environment.

## INTRODUCTION

Genome streamlining provides a fitness advantage to abundant aquatic microorganisms by reducing the resources required to maintain and replicate their genomes (Giovannoni et al., 2005, 2014; Rodríguez-Gijón et al., 2025; Swan et al., 2013). However, genome reduction is constrained by the need to retain functions important for organismal fitness. Consequently, streamlined genomes reflect not only which functions can be lost, but also which functions cannot (Morris et al., 2012). In aquatic microorganisms, streamlining is often associated with reduced physiological flexibility and metabolic specialization (Carini et al., 2013), and the loss of biosynthetic genes that can generate metabolic dependencies (Pacheco-Valenciana, Tausch, et al., 2025) when the corresponding metabolites are obtained from the environment or other community members (Kost et al., 2023). At the same time, genome reduction does not affect all functions uniformly (Rodríguez-Gijón et al., 2025), suggesting that the metabolic capabilities retained in streamlined genomes may provide insight into the functions that remain important for ecological success.

B-vitamin biosynthesis represents an interesting example of the complex relationship between genome size and metabolic potential (Hessler et al., 2023) because of a mismatch between cellular demand and biosynthetic investment: B-vitamins are mostly required in relatively small amounts (Sañudo-Wilhelmy et al., 2014), yet their synthesis often involves complex multi-step pathways that are unevenly distributed across microbial genomes (Gómez-Consarnau et al., 2018; Paerl et al., 2018; Shelton et al., 2019). Moreover, dissolved B-vitamins (B_1_, B_7_ and B_12_) are present at extremely low concentrations throughout large parts of the global oceans (Sañudo-Wilhelmy et al., 2012; Suffridge et al., 2017). This scarcity profoundly constrains the activity, abundance and taxonomic composition of marine microbial communities (Bertrand et al., 2007; Browning et al., 2017; Wienhausen et al., 2022). While genome streamlining generally promotes loss of biosynthetic costly genes in favor of public goods, the scarcity of coenzymes may drive the maintenance of de novo synthesis. Consistent with the genome streamlining hypothesis, complete B-vitamin biosynthetic pathways are less frequently encoded by microorganisms with smaller genomes, particularly among abundant bacteria and archaea (Gregor et al., 2025; Pacheco-Valenciana et al., 2026; Rodríguez-Gijón et al., 2025), yet some lineages contain more genes involved in B-vitamin biosynthesis than expected from genome size alone (Zoccarato et al., 2022), raising the question “why some streamlined lineages retain complex and metabolically costly pathways?”

Despite their small genomes, several ammonia-oxidizing archaea (AOA) such as the members of the genus *Nitrosopumilus* retain pathways for B-vitamin biosynthesis and production (Doxey et al., 2015; Qin et al., 2017). Phylogenomic evidence further suggests that biotin (B_7_) and cobalamin (B_12_) biosynthesis were acquired during the evolutionary transition to AOA as oxygen availability increased, potentially supporting the newly established aerobic metabolism through their roles as cofactors (Ren et al., 2019). Their retention may also have ecological implications in environments where B-vitamin biosynthetic capacity is unevenly distributed. In the Baltic Sea, for example, many bacterioplankton have incomplete thiamine (B_1_), biotin (B_7_), and cobalamin (B_12_) biosynthesis pathways (Pacheco-Valenciana et al., 2026) and instead rely on uptake or salvage strategies that vary across taxa and seasons (Paerl et al., 2018; Pérez-Martínez et al., 2025).

Here we investigated an abundant *Nitrosopumilus* population from the Baltic Sea with a small genome yet high B-vitamin biosynthetic potential. We combined metabolic annotation, population genetics, and phylogenomics to study B-vitamin biosynthesis in this population and across AOA. We also measured B-vitamins in *Nitrosopumilus maritimus* to assess whether biosynthetic potential is reflected in vitamin production. Together, these analyses allowed us to examine the evolutionary and ecological significance of retaining B-vitamin biosynthesis in a streamlined and abundant microbial population.

## MATERIALS & METHODS

### Baltic Sea metagenomes and genome catalog

We used a publicly available genome catalog from the Baltic Sea, consisting of 1603 metagenome-assembled genomes (MAGs) reconstructed from metagenomes of four independent studies spanning different locations and depths across the Baltic Sea (Alneberg et al., 2018, 2020; Larsson et al., 2014; Pacheco-Valenciana, Tausch, et al., 2025). These MAGs were dereplicated into 701 clusters at 95% average nucleotide identity (ANI). For each cluster, the highest-quality MAG (>90% completeness and <5% contamination) was selected as the representative genome and used for downstream analysis. Genome completeness, contamination, and additional genome characteristics were estimated with CheckM (Parks et al., 2015), while taxonomic classification was done using GTDB-Tk (Chaumeil et al., 2020) with release 220 (**Supplementary Data 1**).

### Relative abundance and prevalence estimation

To estimate relative abundances across the Baltic Sea, reads from 112 metagenomic samples (Alneberg et al., 2018, 2020; Larsson et al., 2014; Pacheco-Valenciana, Tausch, et al., 2025) were recruited against the 701 representative genomes using Strobealign (Sahlin, 2022). Resulting BAM files were processed in anvi’o (Eren et al., 2020) by first sorting them with ‘anvi-init-bam’ and subsequently calculating genome coverage across samples with ‘anvi-profile-blitz’. To reduce the influence of weak or partial recruitments, coverage values were only retained when genome detection (coverage breadth) was at least 50%. Relative abundance was then calculated by normalizing each retained genome Q2Q3 coverage value (average coverage after removing the 25% lowest- and 25% highest-coverage nucleotide positions) by the summed retained Q2Q3 coverage of all genomes within the same sample. Prevalence for each genome was defined as the proportion of metagenomes in which it was detected with a relative abundance greater than zero after the detection filtering.

### Genome annotation and pathway completeness estimation

Functional annotation and biosynthetic pathway completeness estimates were performed in anvi’o using KEGG Orthologs derived from the KOfam database (Aramaki et al., 2020), together with annotations from the Pfam protein domain families database (Mistry et al., 2021) and the Clusters of Orthologous Genes (COGs) database (Tatusov, 2000). Biosynthetic pathway completeness was estimated with ‘anvi-estimate-metabolism’ using a user-defined metabolic module description for 18 amino acids and 9 B-vitamins (Pacheco-Valenciana, Tausch, et al., 2025). In order to identify microorganisms with unusually high B-vitamin biosynthetic potential, we calculated a cumulative completeness metric by summing the pathway completeness values for vitamins B_1_, B_7_, and B_12_ across the Baltic Sea genome catalog (**Supplementary Data 1**). A value of three indicated the potential to produce all three B-vitamins de novo, whereas lower values reflected either the capacity to produce at least one B-vitamin or the presence of partial pathways for any or all of the three B-vitamins.

METABOLIC (Zhou et al., 2022) was also used as a complementary tool to identify and interpret key marker genes involved in biogeochemical processes, including carbon fixation and organic carbon and nitrogen metabolism.

### Population genetics analyses

Population genetic analyses focused on the Baltic Sea population represented by the archaeal genome P1994_129_bin89 (ERZ1291182; (Alneberg et al., 2020), classified in GTDB (release 220) as *Nitrosopumilus* sp001437625. We propose the name *Nitrosopumilus supplementum* sp. nov. for the species represented by this genome (**Supplementary Note 1**) and refer to the population as *N. supplementum* throughout the study. The reference genome consisted of 198 contigs and was estimated by CheckM to be 100% complete with 0.97% contamination, while anvi’o estimated 98.68% completion and 6.58% redundancy. Therefore, we used this genome as the reference to examine the *N. supplementum* population across Baltic Sea metagenomes.

We used read recruitment profiles generated in anvi’o (v9) that included single-codon variants (SCVs), using the –profile-SCVs option in the ‘anvi-profilè program. Individual sample profiles were then merged with ‘anvi-mergè, and the focal archaeal genome was extracted for downstream analyses with ‘anvi-split’.

To minimize noise associated with low-coverage metagenomes, only samples with a mean Q2Q3 genome coverage greater than 30× were retained, resulting in 22 metagenomes (**Supplementary Data 2**). Coverage patterns for the 198 contigs from the reference genome were also inspected using ‘anvi-refinè across all metagenomes to assess their consistency. For visualization, **Supplementary Figure 1** shows the 22 samples retained for population genetic analyses. Single-nucleotide variant (SNV) profiles were generated using ‘anvi-gen-variability-profilè, considering only coding regions, requiring a minimum coverage of 30× across all samples and a minimum departure from reference of 10%. Analyses were performed in –quince-mode to retain nucleotide-level variability information suitable for downstream population genetic analyses.

Population differentiation among metagenomes was estimated using fixation indices (FST) calculated from SNV profiles with ‘anvi-gen-fixation-index-matrix’ (Kiefl et al., 2023). The resulting pairwise distance matrix was converted into a hierarchical clustering dendrogram using ‘anvi-matrix-to-newick’ and visualized with ‘anvi-interactivè together with environmental metadata associated with each metagenome.

To investigate whether B-vitamin biosynthesis genes experience distinct selective pressures relative to the rest of the genome, the proportion of non-synonymous to synonymous mutations at the population level (pN/pS) was estimated for individual genes across the 22 metagenomes using the program ‘anvi-get-pn-ps-ratiò (Kiefl et al., 2023). Genes associated with B-vitamin biosynthesis were identified from custom metabolic module annotations corresponding to B_1_ (M00896), B_7_ (M00123), and B_12_ biosynthesis (M00924 and M00122).

### Nitrososphaerales genome phylogeny

A concatenated marker gene phylogeny of the species representative genomes in the *Nitrososphaerales* order was generated based on the genomes included in the GlobDB (Speth et al., 2025) (release 226). Species representative genomes of the orders *Conexivisphaerales*, *Geothermarchaeales*, JACIWG01, and PSMU01, which are all part of the *Nitrososphaeria* class, were used as an outgroup. Only the GlobDB species representatives with an estimated genome completeness of >70% were included in this analysis. The phylogeny was generated based on the anvi’o (Eren et al., 2020) contigs databases available from the GlobDB website (https://globdb.org/). Amino acid sequences of the 76 archaeal marker genes with HMMs available in anvi’o (https://github.com/merenlab/anvio/tree/master/anvio/data/hmm/Archaea_76) were exported using the program ‘anvi-get-sequences-for-hmm-hits’. Outliers in the exported HMM matches were removed using the ‘filter’ functionality of the amino acid sequence toolkit (https://github.com/dspeth/aastk). The filtered datasets were then aligned using Muscle5 (Edgar, 2022) (v5.3), and alignments were trimmed using TrimAl (Capella-Gutiérrez et al., 2009) (v1.5.1) with the -gappyout option. Trimmed alignments were concatenated using the catfasta2phyml script (https://github.com/nylander/catfasta2phyml). A phylogeny of the concatenated alignment was generated using FastTree (Price et al., 2010) (v2.2.0).

Presence or absence of genes involved in B vitamin biosynthesis in the genomes included in the phylogeny was assessed based on KEGG annotations included in the GlobDB anvi’o databases. These annotations were generated using the adaptive adjustment of significance thresholds integrated in anvi’o (Kananen et al., 2025).

### *Nitrosopumilus maritimus* SCM1 growth and LC-MS analysis of intra- and extracellular B-vitamins

*Nitrosopumilus maritimus* SCM1 was grown in 200 mL of modified synthetic crenarchaeota medium (SCM1; Martens-Habbena et al., 2009) without the addition of vitamins or substrates. Cultures were grown in triplicate at 28°C in the dark without shaking. Growth was monitored through nitrite production using sulfanilamide/NED reagent, with absorbance measured at 545 nm and a 0.5 mM nitrite solution used as a reagent control. During the late exponential growth, cultures were filtered through 0.1-µm polyethersulfone filters. Filters containing the cells and the corresponding exometabolomes were stored at −20°C until further analysis. Samples were also collected at the same sampling point for cell enumeration using flow cytometry (BD Accuri plus, BD Biosciences, Franklin Lakes, NJ, USA).

Extracellular B-vitamins and precursors were extracted from the archaeal exometabolome by solid-phase extraction using Bond Elut PPL (1 g; Agilent, Santa Clara, CA, USA). Before extraction, the pH of the samples was adjusted to 6 using 37% hydrochloric acid, and the solid-phase extraction columns were conditioned with 10 mL of methanol. Extracted exometabolomes were eluted with ULC/MS-grade methanol (Biosolve, Dieuze, France), evaporated by nitrogen fumigation at 28°C, and redissolved in 200 µL of ultrapure water produced using an arium® pro ultrapure water system (Sartorius, Göttingen, Germany), as described elsewhere (Bruns et al., 2023).

Intracellular B-vitamins and precursors were extracted from filters through three rounds of bead beating using a Mini-Beadbeater-24 (Clifton, New York, USA), as described elsewhere (Cakić et al., 2021). Standards (Sigma-Aldrich, Steinheim, Germany) were used to evaluate the recovery rate. Concentrated intracellular and extracellular samples were analyzed by LC-MS as described elsewhere (Bruns et al., 2022), using an Ultimate 3000 HPLC coupled to a SQ Quantum UltraAM triple quadrupole mass spectrometer (both Thermo Fisher Scientific) on a Kinetex Evo C18 column (100 × 2.1 mm, 2.6 µm pore size; Phenomenex, Torrance, CA, USA). Ten mM ammonium formate at pH 6.0 (A) and acetonitrile (B) were used with 0–13 min 100–75% A; 13–15min 75–0% A; 15–19 min 0% A; 19–21min 0–100% A; 21–26 min 100% as solvent gradient. Metabolite concentrations were expressed as molecules per archaeal cell.

### Statistical analyses

For statistical analyses and visualization, we used R and RStudio (v.4.40). To test differences in pN/pS between B-vitamin biosynthesis genes and the remaining genes in the genome, we used the Mann-Whitney U test (Mann & Whitney, 1947). The associations between selective pressure and environmental variables were evaluated using Spearman’s rank correlation (Spearman, 1987).

## RESULTS

### An abundant *Nitrosopumilus* population with a streamlined genome retains high B-vitamin biosynthetic potential

We compared estimated genome size and the cumulative completeness of biosynthetic pathways for vitamins B_1_, B_7_, and B_12_ across all representative genomes recovered from the Baltic Sea catalog (Pacheco-Valenciana, Garcia, et al., 2025; Pacheco-Valenciana, Tausch, et al., 2025). Among these, the *N. supplementum* genome stood out by combining a small genome size with high cumulative completeness of B-vitamin biosynthetic pathways (**Figure 1**). It had an estimated genome size of 1.4 Mbp (**Figure 1A**) and a cumulative pathway completeness of 2.4 for B_1_, B_7_, and B_12_ (**Figure 1C**). In addition to its unusual combination of genomic characteristics, *N. supplementum* was prevalent in 49% of the analyzed Baltic Sea metagenomes, with an average relative abundance of 1.46% (**Figure 1B**).

**Figure 1.**
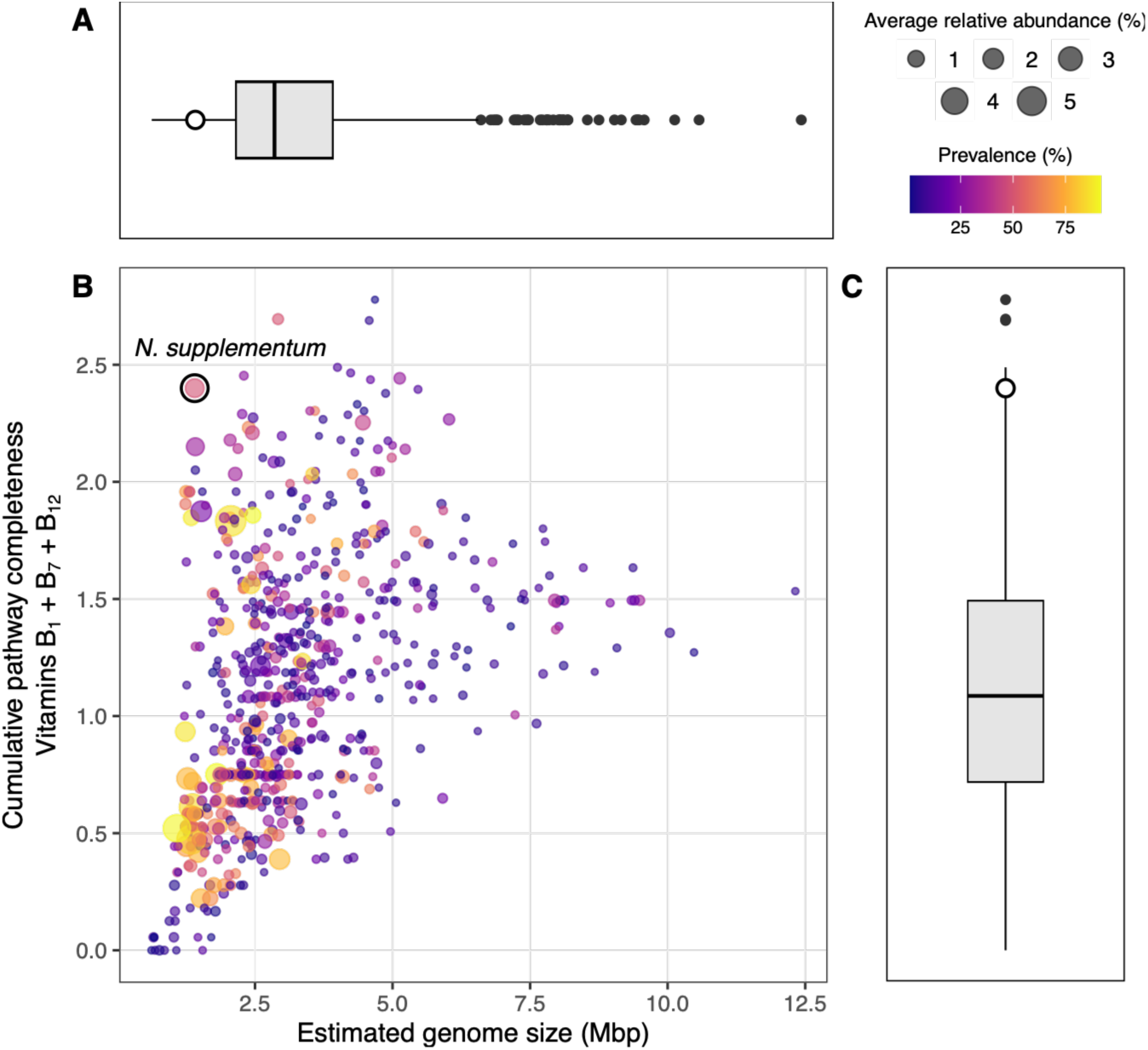
Estimated genome size, B-vitamin biosynthetic potential, and ecological distribution across the Baltic Sea genome catalog. (A) Boxplot showing the distribution of estimated genome sizes (Mbp) across all genomes in the Baltic Sea catalog. (B) Scatter plot showing the relationship between estimated genome size (Mbp) and cumulative pathway completeness for three B vitamins (B1, B7, and B12) across all genomes. Cumulative pathway completeness represents the sum of module completeness values for the three vitamins (range: 0–3). Each point corresponds to a genome. Point size indicates the average relative abundance (%) across all samples, and point color represents prevalence (%). (C) Boxplot showing the distribution of cumulative pathway completeness for vitamins B1, B7, and B12 across all genomes, using the same definition as in panel B. The archaeal genome of interest, *N. supplementum*, is highlighted by a black-outlined circle in all panels and labeled in panel B.

The genome-scale metabolic reconstruction revealed a chemolithoautotrophic metabolism characteristic of marine AOA, including pathways associated with ammonia oxidation and carbon fixation, together with genes for ammonium and urea transport and the urease complex (**Supplementary Figure 2**). Central carbon metabolism included gluconeogenesis and partial TCA and pentose phosphate pathways, while the genome also encoded several ABC transport systems involved in the uptake of metals, nutrients, and small organic compounds. The genome also retained substantial B-vitamin biosynthetic potential. More specifically, this archaeal genome encoded a complete de novo vitamin B_1_ (thiamine) biosynthesis pathway, including the *thi4* gene associated with thiazole biosynthesis, *thiC* and *thiD* associated with the HMP biosynthesis (pyrimidine branch), *thiN*, which is responsible for coupling both moieties, and t*hiL*, which catalyzes the final step. For B_7_ (biotin), the genome encoded all genes associated with KEGG module M00123 but we did not detect genes associated with the annotated pathways for pimeloyl-ACP and pimeloyl-CoA biosynthesis, the primary precursors for B_7_. As for B_12_ (cobalamin), the genome has an almost complete anaerobic corrin ring pathway, lacking only *cbiJ*, and an incomplete nucleotide loop assembly pathway. Genes associated with corrinoid remodeling are present, along with the *bluB* gene, responsible for the synthesis of the lower ligand 5,6-dimethylbenzimidazole (DMB). Given its high prevalence and unusual combination of genome streamlining and B-vitamin biosynthetic potential, we next investigated the population structure of *N. supplementum* across the Baltic Sea.

### The *N. supplementum* population shows genetic differentiation across the Baltic Sea

Read recruitment detected the *N. supplementum* population in 55 of the 112 analyzed metagenomes, with relative abundances ranging from less than 0.01% to 30.11%. Of these, 22 metagenomes had more than 30× coverage of the *N. supplementum* population. These metagenomes spanned broad geographic regions of the Baltic Sea, including northern, central, southern, and those near the Atlantic entrance (**Supplementary Data 2**).

We calculated SNV (single nucleotide variant) profiles and estimated the genetic differentiation between samples using pairwise fixation indices (FST). Hierarchical clustering based on the resulting FST values separated the metagenomes into five major groups (G1–5) (**Figure 2**). These groups showed differences in geographic distribution and environmental conditions, including salinity, temperature, depth, and season. Irrespective of their location, all surface samples formed one major cluster (G2), with a subgroup of samples near the Gotland region, and another subgroup in more southern coastal environments. A second major group (G1) included samples with salinity between 7.3 and 12 PSU and was further subdivided by depth and sampling season. The most distinct high-salinity group (G4) consisted of two samples (ERR2206752 and ERR2206753), collected near the Atlantic entrance. These samples also showed the highest relative abundance of this *N. supplementum,* reaching 15.06% and 30.11%, respectively.

**Figure 2.**
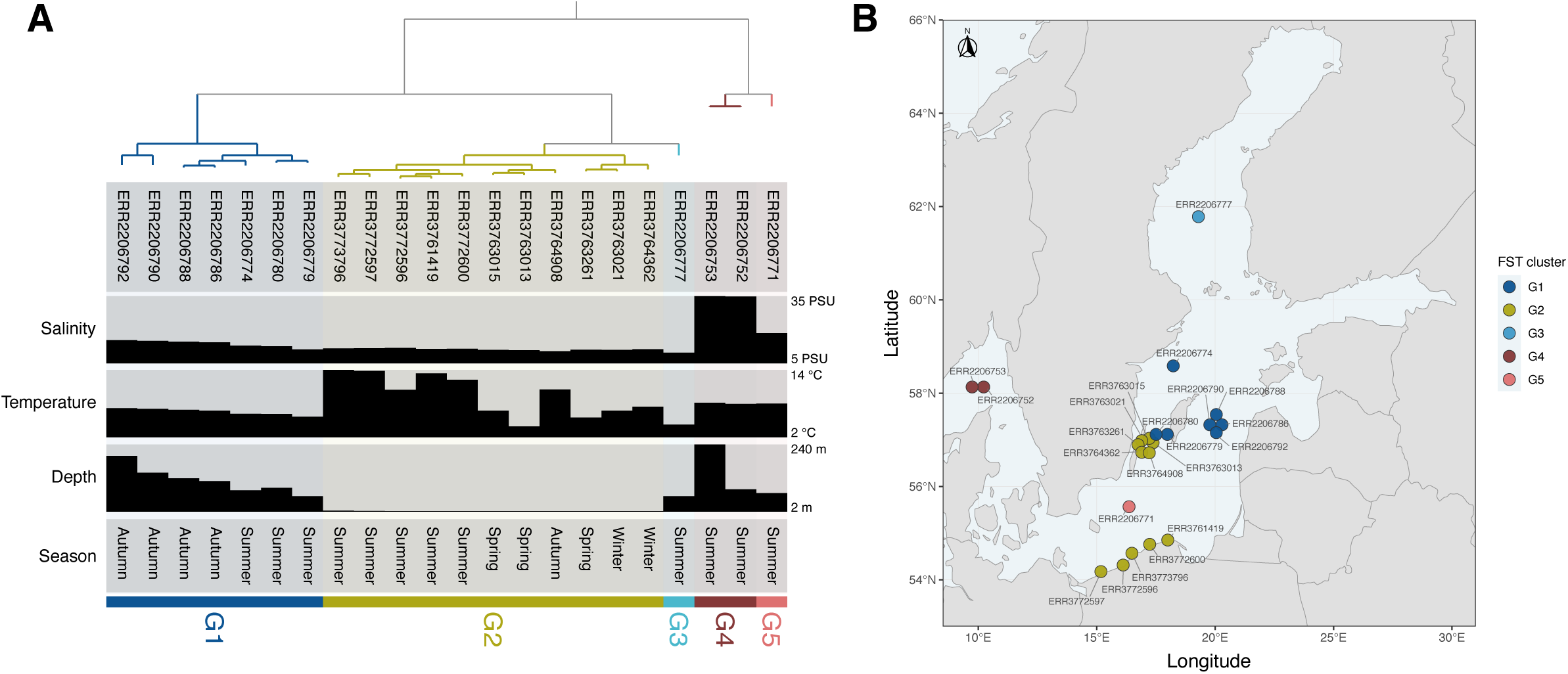
Population structure and geographic distribution of the *Nitrosopumilus* population across the Baltic Sea. (A) Hierarchical clustering of metagenomes based on population-level genetic variation within the *Nitrosopumilus* population. Samples were grouped according to genomic similarity across sites with sufficient coverage for population genetic analyses. Colored branches and labels indicate the major population clusters (G1–G5). Environmental metadata associated with each metagenome, including season, salinity (PSU), temperature (°C), and sampling depth (m), are shown. Grayscale bars represent the observed values of continuous environmental variables, with their ranges indicated on the right side of the plot. (B) Geographic distribution of the archaeal population identified in panel A across Baltic Sea sampling stations. Each point represents a metagenome, colored by its assigned population cluster. Points at co-located sampling stations are slightly offset for visibility.

The geographic distribution of these population groups also revealed spatial structure across the Baltic Sea (**Figure 2B**). Samples belonging to the same genomic clusters generally occurred in similar geographic regions, suggesting that population differentiation within this archaeal population is associated with environmental variation across the Baltic Sea, particularly the strong salinity gradient, as has been observed earlier for this population in an analysis encompassing only surface samples (Sjöqvist et al., 2021). These patterns show that the *Nitrosopumilus* population is not only widespread and abundant, but also genetically structured across contrasting Baltic Sea conditions. This provided a basis to examine whether the B-vitamin biosynthesis genes retained in this streamlined genome show evidence of selective constraint within this natural population.

### B-vitamin biosynthesis genes in *N. supplementum* show indications of strong purifying selection

Of the 25 genes initially annotated as part of B-vitamin biosynthetic modules in this genome, including genes associated with B_1_ (M00896), B_7_ (M00123), and B_12_ (M00924 and M00122) (**Supplementary Data 3**), we excluded two genes from further analyses because they are not specific to B-vitamin biosynthesis. These included uroporphyrin-III C-methyltransferase (*cysG*/*cobA*, anvi’o gene call 1401948) and precorrin-2 dehydrogenase (*sirC*, anvi’o gene call 1402527), as both of these enzymes catalyze early steps in tetrapyrrole metabolism that are shared between siroheme and cobalamin biosynthesis. We also excluded dethiobiotin synthetase (*bioD*, anvi’o gene call 1402037) because sufficient sequencing coverage was detected in only 2 of the 22 metagenomes, producing unreliable pN/pS estimates. Of the remaining 22 B-vitamin associated genes, 19 had at least one usable pN/pS value across metagenomes, while three genes (anvi’o gene calls 1402096, 1402313, and 1402638) lacked data entirely due to insufficient coverage in all samples.

Overall, B-vitamin biosynthesis genes exhibited lower pN/pS ratios than the genomic background, suggesting that they experience stronger purifying selection. The mean per-gene pN/pS ratio for the 19 B-vitamin genes was 0.057, compared to 0.206 for the remaining 1,155 genes in the genome, indicating a nearly four-fold difference (**Figure 3B**, **Supplementary Data 4**). While this difference did not reach statistical significance (Mann-Whitney U test, p = 0.102), the average B-vitamin gene experienced stronger purifying selection than 53% to 83% of genes in this genome across metagenomes. The relative strength of this pattern also varied geographically. The two metagenomes sampled closest to the Atlantic Ocean entrance into the Baltic Sea (ERR2206752 and ERR2206753; Skagerrak Strait, ∼10°E) showed the weakest relative selective signal on B-vitamin genes (53% and 55%, respectively), while several metagenomes from the interior Baltic exceeded the 70th percentile (**Figure 3A**). However, this trend was not statistically significant (Spearman rho = 0.32, p = 0.14 for longitude).

**Figure 3.**
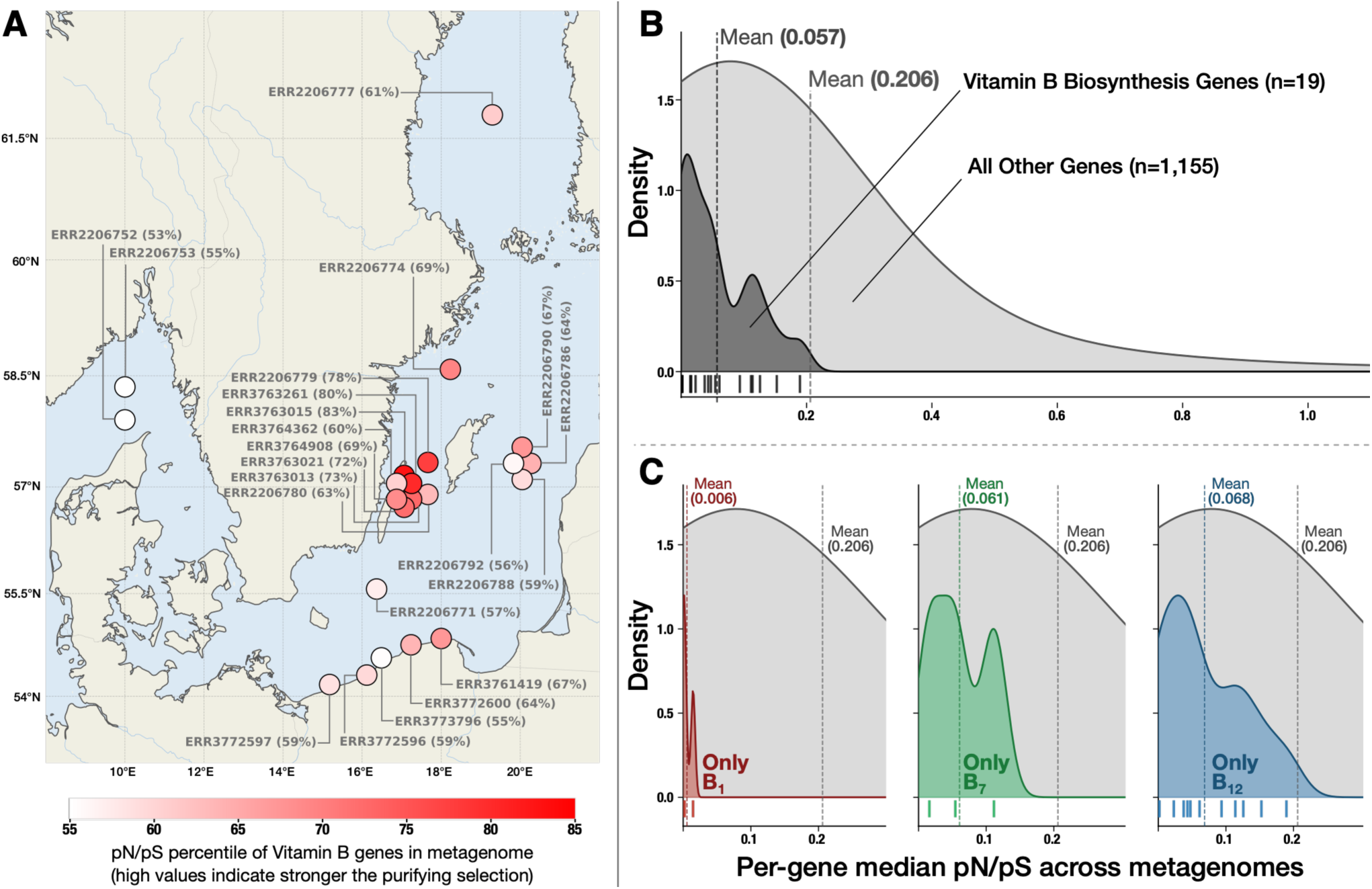
Selective pressure on vitamin B biosynthesis genes in the Baltic Sea archaeon. (A) Map shows the geographic distribution of selective pressure on vitamin B genes across 22 Baltic Sea metagenomes. Each point represents a metagenome, colored on a continuous scale by the average percentile rank of vitamin B genes relative to all genes in that sample. Percentages show the average percentile rank for each sample. Points at co-located sampling stations are offset slightly for visibility. (B) Distribution of per-gene median pN/pS values for all B-vitamin biosynthesis genes combined compared to all other genes in the genome. Each gene’s median pN/pS was computed across all metagenomes in which it had sufficient coverage. Black tick marks along the x-axis show individual vitamin B gene values. Dashed vertical lines indicate the mean of each distribution. (C) Density plot showing the same comparison broken down by individual vitamin B pathways.

Selective pressure also varied among individual genes within the three B-vitamin biosynthesis pathways (**Figure 3C**). B_1_ biosynthesis genes were under the most consistent and strongest purifying selection, with *thiC* (phosphomethylpyrimidine synthase) having a median pN/pS value of zero across metagenomes, placing it at the 100th percentile genome-wide, while *thiL* (thiamine-monophosphate kinase) ranked at the 85th percentile. The cobalamin biosynthesis pathway, in contrast, showed an internal gradient. The late-pathway enzymes responsible for amidation (*cbiA*, 61st percentile; *cbiP*, 100th percentile) and adenosylation (*btuR*, 85th percentile) were under substantially stronger constraint than mid-pathway ring methyltransferases such as c*biF* (36th percentile), *cbiG* (28th percentile), and *cbiT* (33rd percentile).

Interpretation of these patterns should be done cautiously due to differences in sequencing coverage among genes. Genes with stronger signatures of purifying selection were generally represented across 20–22 metagenomes, whereas several genes with weaker selective signals were only detected in 5–8 metagenomes. Among the six corrin ring methyltransferases, the two genes with complete metagenome representation (*cbiL* and *cbiH*) ranked within the top tier of purifying selection (100th and 70th percentile, respectively), suggesting that part of the observed variability among pathway genes may reflect differences in coverage across metagenomes.

Further evidence for the functional importance of B-vitamin biosynthesis in this population comes from the vitamin-dependent enzymes encoded in this genome. For B_1_, the genome encoded two TPP-dependent enzymes, acetolactate synthase, involved in branched-chain amino acid biosynthesis, and transketolase, involved in central carbon metabolism through the pentose phosphate pathway. The presence of acetolactate synthase is particularly significant because the valine and isoleucine biosynthesis pathways were complete in this genome, directly linking B_1_ availability to these retained amino acid biosynthetic functions. For B_12_, the genome encoded the cobalamin-dependent methionine synthase (*metH*), whereas the B_12_-independent alternative (*metE*) was absent, indicating that the detected methionine synthase route depends on cobalamin availability. These annotations provide additional genomic information for interpreting the selective constraint observed on B-vitamin biosynthesis genes.

Despite some limitations, the retention and selective constraint of B-vitamin biosynthesis genes in *N. supplementum* raised two broader questions: whether these functions are conserved across related archaea and whether this biosynthetic potential corresponds to measurable vitamin production.

### B-vitamin biosynthesis genes are broadly conserved across AOA

To place the B-vitamin biosynthesis potential of *N. supplementum* in a broader phylogenetic context, we examined the distribution of genes associated with B_1_, B_7_, and B_12_ biosynthesis across 1,077 archaeal species representatives (**Figure 4**). The dataset comprised 1,010 representatives within the order *Nitrososphaerales*, 838 of which are AOA, along with 67 non-AOA representatives from four other orders within the *Nitrososphaeria* class used as phylogenetic outgroups, namely *Conexivisphaerales*, *Geothermarchaeales*, *JACIWG01*, and *PSMU01* (**Supplementary Data 5**). Within the *Nitrososphaerales* order, *Nitrosopumilus* was the most represented genus, with 145 species representatives. Across the phylogeny, AOA representatives encoded many of the genes associated with B_1_, B_7_, and B_12_ biosynthesis, showing broad conservation within the *Nitrososphaerales* order (Ren et al., 2019). B_1_ biosynthesis showed a particularly consistent pattern across AOA in our study. The first step of the archaeal-specific M00896 pathway, represented by K22699 (*thi4*), was broadly conserved across AOA, together with several genes involved in the subsequent synthesis of thiamine. B_7_ biosynthesis was also broadly represented, although conservation differed among individual steps of the pathway. K01935 (*bioD*) and K01012 (*bioB*), which catalyze the final steps leading to biotin, were particularly widespread across AOA and largely absent from the outgroup, whereas K00652 (*bioF*) showed a more variable distribution. B_12_ biosynthesis showed the most complex distribution. Genes associated with anaerobic corrin ring biosynthesis were broadly conserved across much of the AOA phylogeny, whereas genes associated with nucleotide-loop assembly showed a more uneven distribution, with some steps only rarely detected. In comparison, the outgroup generally showed a lower presence of genes associated with all three B-vitamin pathways. The broad distribution of these genes across AOA indicates that B-vitamin biosynthetic capacity is a conserved feature of this archaeal lineage rather than a unique characteristic of *N. supplementum*.

**Figure 4.**
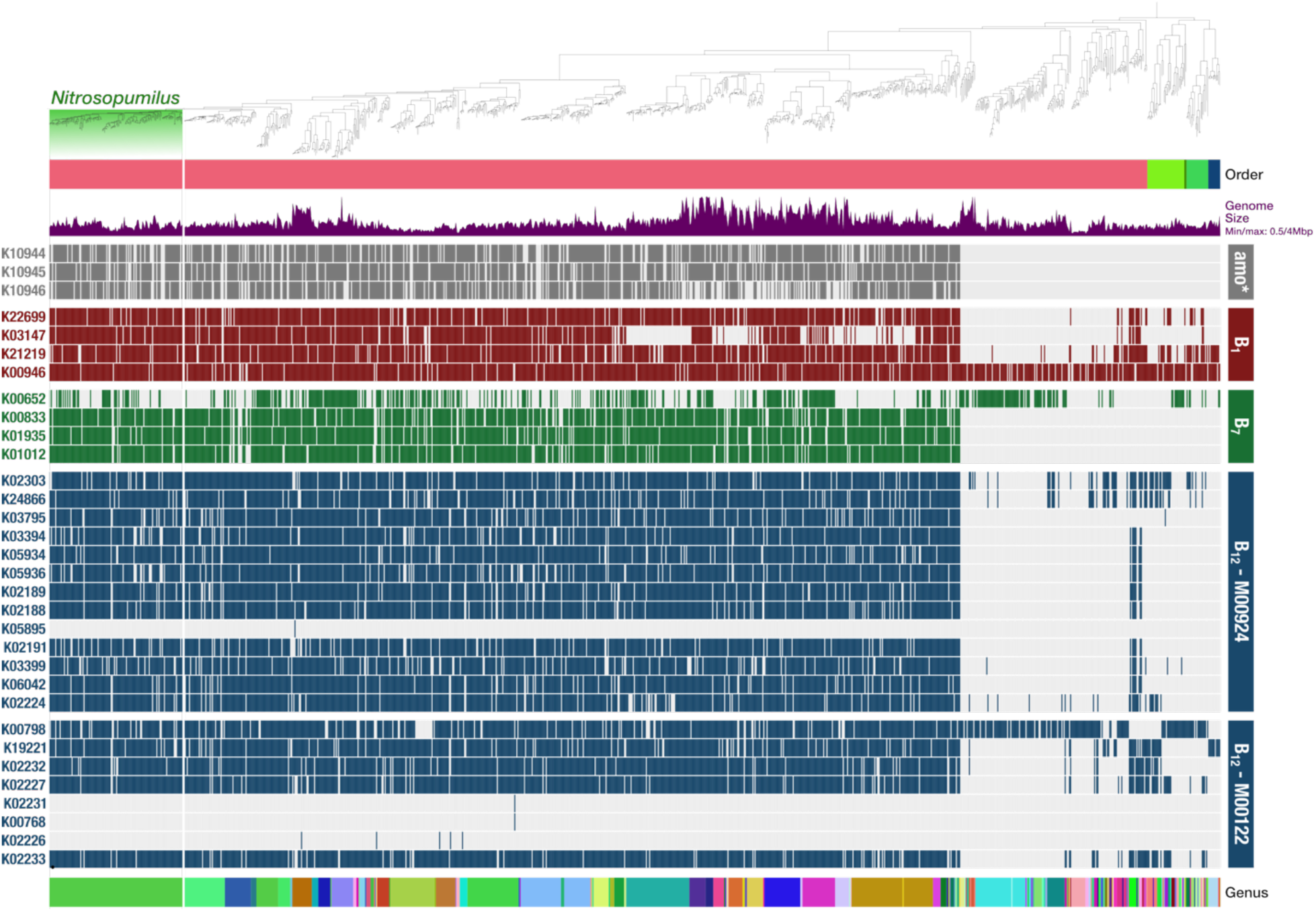
Phylogenetic distribution of B-vitamin biosynthesis genes across archaeal genomes. Phylogenomic tree of 1,077 archaeal species representatives with annotations for taxonomic order and genome size. The heatmap shows the presence or absence of KEGG Orthologs (KOs) associated with ammonia oxidation and vitamin B_1_, B_7_, and B_12_ biosynthesis across genomes, ordered by phylogeny. Ammonia oxidation genes (amo* = *amoABC*) are shown in grey, B_1_ biosynthesis genes (M00896) in red, B_7_ biosynthesis genes (M00123) in green, and B_12_ genes belonging to the anaerobic corrin-ring biosynthesis (M00924) and nucleotide-loop assembly (M00122) modules in blue. Filled cells indicate gene presence, whereas unfilled cells indicate that the corresponding gene was not detected. The *Nitrosopumilus* clade is highlighted in the phylogeny using the same color as the genus annotation shown at the bottom.

### B-vitamins and their precursors are detected in intracellular and extracellular fractions of *Nitrosopumilus maritimus*

Although an isolate corresponding to *N. supplementum* is not available, the cultured marine AOA *N. maritimus* allowed us to examine whether the biosynthetic potential observed across AOA corresponds to measurable intracellular and extracellular vitamins in this closely related species.

Among the vitamins examined here, B_1_ showed the highest mean concentration in both fractions, with substantially higher intracellular than extracellular levels. B_7_ was only detected in the extracellular fraction and showed a higher mean concentration than the detected extracellular B_12_ forms. Cyanocobalamin (CB_12_) and adenosylcobalamin (AB_12_) were detected in both fractions, with adenosylcobalamin more abundant than cyanocobalamin in both cases, while methylcobalamin (MB_12_) was only detected intracellularly at lower concentrations than the previous B_12_ forms. Several vitamin precursors were also detected extracellularly, including HMP, HET, desthiobiotin (dB_7_), and the lower-ligand compound DMB, with DMB showing a particularly high mean extracellular concentration compared with the measured B-vitamins (**Table 1**). Thus, multiple B-vitamins and vitamin precursors were detectable in *N. maritimus*, including several compounds present in the extracellular fraction.

**Table 1:** Intracellular and extracellular concentrations of B-vitamins and precursors extracted from *Nitrosopumilus maritimus* SCM1.

| Vitamin/Precursor | Intracellular vitamin molecules per cell | Extracellular vitamin molecules per cell |
| --- | --- | --- |
| B<sub>1</sub> | $8.21 \times 10^6 \pm 8.51 \times 10^5$ | $3.12 \times 10^5 \pm 2.91 \times 10^5$ |
| B<sub>2</sub> | $1.93 \times 10^6 \pm 3.10 \times 10^5$ | $6.93 \times 10^5 \pm 1.58 \times 10^5$ |
| B<sub>3</sub> | $5.83 \times 10^5 \pm 2.55 \times 10^5$ | ND |
| B<sub>5</sub> | $4.14 \times 10^6 \pm 1.32 \times 10^6$ | ND |
| B<sub>6</sub> | $1.23 \times 10^4 \pm 2.06 \times 10^3$ | $1.44 \times 10^4 \pm 3.79 \times 10^3$ |
| B<sub>7</sub> | ND | $8.34 \times 10^4 \pm 2.39 \times 10^4$ |
| B<sub>9</sub> | ND | ND |
| CB<sub>12</sub> | $1.57 \times 10^5 \pm 3.67 \times 10^4$ | $1.30 \times 10^4 \pm 5.44 \times 10^3$ |
| AB<sub>12</sub> | $5.35 \times 10^5 \pm 1.10 \times 10^5$ | $3.64 \times 10^4 \pm 3.31 \times 10^4$ |
| MB<sub>12</sub> | $5.47 \times 10^3 \pm 3.04 \times 10^3$ | ND |
| HB<sub>12</sub> | ND | ND |
| HMP | ND | $1.81 \times 10^4 \pm 1.86 \times 10^4$ |
| HET | ND | $2.31 \times 10^4 \pm 1.21 \times 10^4$ |
| dB<sub>7</sub> | ND | $1.46 \times 10^5 \pm 4.20 \times 10^4$ |
| Cbi | ND | ND |
| DMB | ND | $4.13 \times 10^6 \pm 4.09 \times 10^6$ |
| α-Rib | $8.44 \times 10^4 \pm 9.28 \times 10^3$ | $9.81 \times 10^5 \pm 5.00 \times 10^5$ |
| Adn | $3.20 \times 10^7 \pm 8.31 \times 10^6$ | $8.49 \times 10^7 \pm 2.07 \times 10^7$ |

\*CB<sub>12</sub>, Cyanocobalamin; AB<sub>12</sub>, Adenosylcobalamin; MB<sub>12</sub>, Methylcobalamin; HB<sub>12</sub>, Hydroxycobalamin; HMP, 4-Amino-5-hydroxymethyl-2-methylpyrimidine; HET, 5-(2-Hydroxyethyl)-4-methylthiazole; dB<sub>7</sub>, Desthiobiotin; Cbi, Cobinamide; DMB, 5,6-Dimethylbenzimidazole; $\alpha$ -Rib, $\alpha$ -Ribazole; Adn, Adenine; ND, not detected.

## DISCUSSION

The abundant *Nitrosopumilus supplementum* population investigated in the Baltic Sea retains substantial biosynthetic capacity for vitamins B_1_, B_7_, and B_12_ despite its streamlined genome. At the same time, many of the metabolic traits identified in this genome are also commonly found in ammonia-oxidizing archaea (AOA), including the amoABC complex associated with ammonia oxidation (Könneke et al., 2005; Walker et al., 2010), carbon fixation through the 3HP/4HB cycle (Könneke et al., 2014), and biosynthetic pathways for several B-vitamins (Bergauer et al., 2025; Doxey et al., 2015). Our phylogenomic analysis is consistent with previous evidence suggesting that biotin (B_7_) and cobalamin (B_12_) biosynthesis were acquired together with ammonia oxidation as AOA evolved in increasingly oxygenated environments (Ren et al., 2019).

The combination of a streamlined genome, broad Baltic Sea distribution, and retention of several B-vitamin biosynthetic pathways provides an interesting ecological and evolutionary perspective within the Baltic Sea microbial community. This particular case suggests that genome streamlining does not always result in the loss of complex biosynthetic functions. Instead, some streamlined genomes may retain metabolic pathways that remain important for fitness under specific community or environmental contexts, while other functions become dispensable through metabolite exchange and ecological interactions (Mas et al., 2016; Morris et al., 2012). The availability of B-vitamins in marine ecosystems, for example, could have a significant impact on the loss of costly and complex biosynthetic genes (Sañudo-Wilhelmy et al., 2012; Suffridge et al., 2017). Notably, the functions retained by *Nitrosopumilus supplementum* included B-vitamin biosynthetic pathways that are frequently incomplete in other Baltic Sea genomes (Pacheco-Valenciana, Tausch, et al., 2025; Paerl et al., 2018).

Population structure added another layer to this pattern. The *Nitrosopumilus* population was not only widespread and locally abundant but also genetically differentiated across the Baltic Sea, with population structure corresponding broadly to geographic and environmental variation. A recent genome-resolved study identified *Ca. Nitrosopumilus balticolittoralis* as a close relative of *N. supplementum* (Kim et al., 2026). Despite their close relationship, the two populations showed different geographic distributions. *Ca. N. balticolittoralis* predominated in western Baltic surface waters, whereas our population was more abundant in central and eastern surface waters. Environmental heterogeneity has previously been associated with genomic differentiation and signatures of local adaptation in different Baltic Sea microbial populations, including this *Nitrosopumilus* population (Sjöqvist et al., 2021). Our analysis extends this previous observation across a broader set of samples spanning both surface and deeper waters.

One of the main signals identified in this study is the low pN/pS ratios associated with B-vitamin biosynthesis genes. These genes generally showed lower pN/pS values than the rest of the genome, which is consistent with strong purifying selection (Hughes, 2007). This pattern suggests that amino-acid-changing variation in these genes is less tolerated, supporting the idea that these pathways are important for this archaeon. Recent studies of SAR11 populations further suggest that metagenomic variation across natural populations can reveal environmentally relevant selective pressures acting on specific metabolic genes and pathways (Kiefl et al., 2023), while also revealing habitat-specificity in abundant bacterial populations from the ocean (Tucker et al., 2025). In this archaeon, it was particularly striking that several B_1_ and B_12_ biosynthesis genes, including *thiC* and *cbiP*, exhibited the strongest selective constraints, with median pN/pS values of 0 across metagenomes. However, not all B-vitamin biosynthesis genes could be evaluated in the same way. In particular, the inconsistent coverage of *bioD* relative to the rest of the genome suggests that this locus may be variably represented across the *N. supplementum* population. Read recruitment analyses have similarly shown that genes that do not consistently co-occur with an otherwise detected population can represent accessory genomic content present only in a fraction of the population across specific environments (Delmont & Eren, 2018).

In the case of B_1_, the evidence of strong purifying selection is particularly interesting because complete de novo biosynthesis appears to be relatively uncommon across the Baltic Sea genome catalog (Pacheco-Valenciana et al., 2026). Widespread B_1_ auxotrophy among Baltic bacterioplankton and temporal changes in B_1_ uptake and salvage strategies indicate potential reliance on externally available B_1_ or its precursors at the community level (Paerl et al., 2018). More recent metatranscriptomic analyses showed that B_1_ synthesis, salvage, and uptake are seasonally partitioned among Baltic microbial populations (Pérez-Martínez et al., 2025). Against this background, complete B_1_ biosynthesis under strong selective constraint in *N. supplementum* is particularly notable. The similarly strong constraint on *cbiP* suggests that this pattern is not restricted to B_1_ but also extends to components of B_12_ biosynthesis. The presence of B-vitamin-dependent metabolic functions, including the B_12_-dependent methionine synthase *metH* in the absence of the B_12_-independent alternative *metE*, provides a direct potential benefit of retaining these biosynthetic capabilities for *N. supplementum*.

This raises a broader question of whether the retention of these biosynthetic pathways benefits primarily the archaeon itself or also the surrounding microbial community. This distinction is important because selection acts most directly on the fitness of the organism carrying these genes, whereas any benefit to neighboring microorganisms would arise as an ecological consequence if vitamins or their precursors become externally available. The two possibilities are not mutually exclusive. A biosynthetic pathway is maintained because it is advantageous to the archaeon and could simultaneously increase the availability of compounds required by organisms with incomplete pathways. In this way, the retention of B-vitamin biosynthesis may connect the metabolic requirements of an individual population with metabolic dependencies across the surrounding community. Whether this occurs in the Baltic Sea population ultimately depends on whether the B-vitamins are already present in sufficient concentrations, which is currently unknown, and under which conditions these coenzymes would be released.

Our measurements in the cultured relative *N. maritimus* SCM1 provide initial support for the possibility that vitamins synthesized by *Nitrosopumilus* can become extracellularly available. While previous cultivation studies showed that *Nitrosopumilus* species produce several B-vitamins, including B_1_ and B_12_ (Qin et al., 2017), here we detected B_1_, B_7_, and several forms of B_12_ and related precursors in the extracellular fraction of *N. maritimus* SCM1. This is consistent with previous reports (Bayer et al., 2019, 2024; Law et al., 2021). Even though our approach is based on cultures and we cannot draw conclusions about the active release of the B-vitamins into the environment, our data set does show that the coenzymes can most likely become available through natural processes such as cell leakage or viral lysis (Sultana et al., 2025). Nevertheless, our results establish that B-vitamins and their precursors synthesized by a close relative of *N. supplementum* can occur extracellularly, providing a potential route through which retained biosynthetic functions in streamlined AOA could contribute to metabolite availability in microbial communities.

More broadly, our findings suggest that metabolic dependencies in natural communities may be understood not only through the biosynthetic pathways that streamlined populations lose, but also through those that they retain.

## ACKNOWLEDGMENTS

We thank Dr. Martin Könneke for providing the strain of *Nitrosopumilus maritimus* SCM1 for the vitamin measurements.

## AUTHOR CONTRIBUTIONS

SLG conceived and supervised the study. AP-V, SLG, AME, and GW refined the research idea. AP-V performed the abundance, prevalence and metabolic potential analysis and wrote the first draft of the manuscript. DRS provided the phylogenetic tree and associated metadata. AFA assisted with the reference genome information. AME conducted the population genetic analysis and visualization. SB and HW performed the B-vitamin analyses and data evaluation. All authors contributed to data interpretation, literature review, and manuscript revisions.

## DATA AVAILABILITY

The *N. supplementum* reference genome P1994_129_bin89 is available in ENA under BioSample SAMEA6483089 and accession ERZ1291182. Accession IDs and associated metadata for the Baltic Sea metagenomes used in this study are provided in **Supplementary Data 2**, and accession IDs for the archaeal genomes included in the phylogenomic analysis are provided in **Supplementary Data 5**. A reproducible bioinformatics workflow for the population genetics and molecular evolution analyses of the Baltic Sea archaeon is available at https://merenlab.org/data/baltic-sea-archaeon/.

## SUPPLEMENTARY FIGURES

**Supplementary Figure 1.**
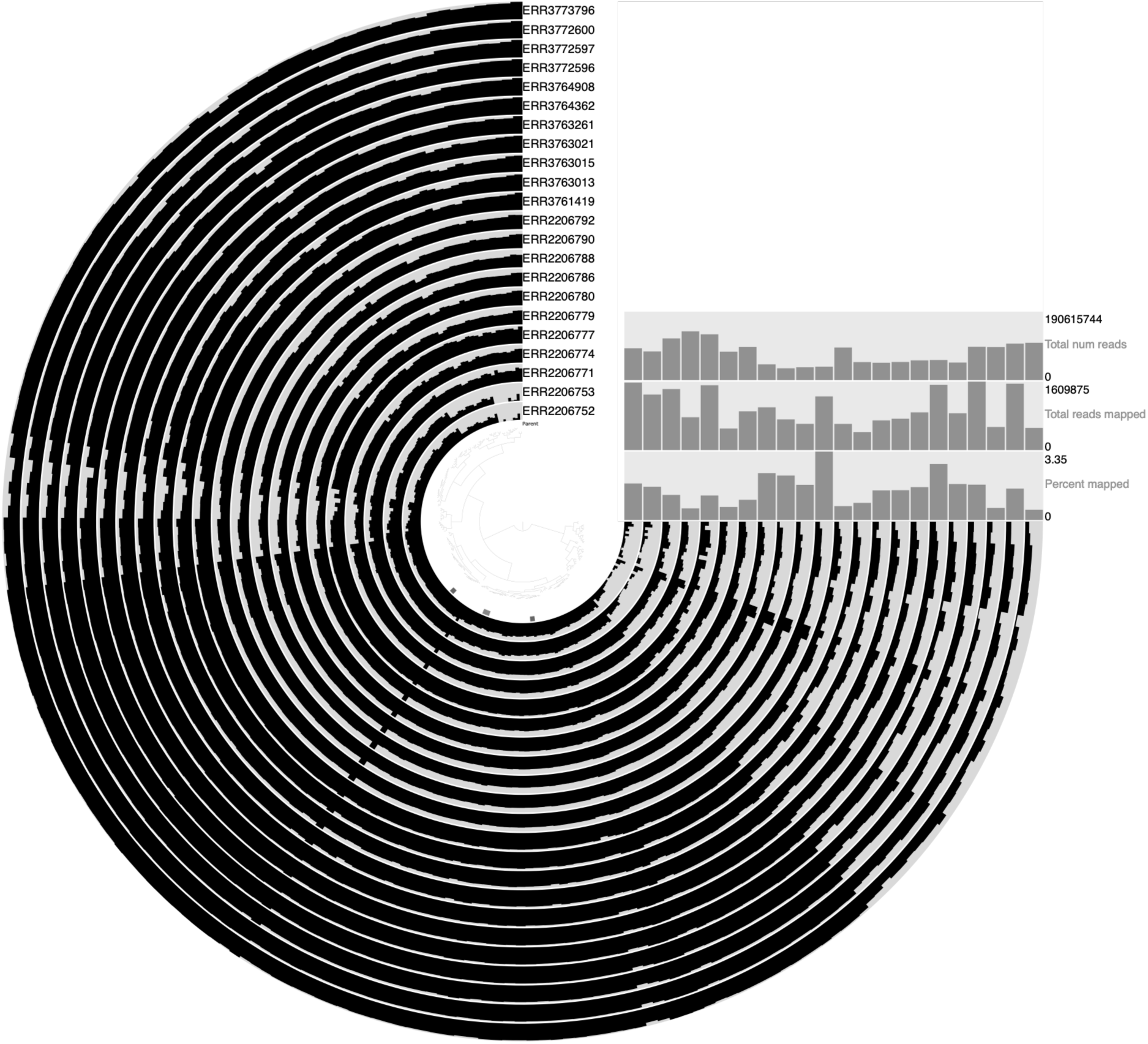
Coverage profiles across the *N. supplementum* reference genome. Mean Q2Q3 coverage profiles for the reference genome P1994_129_bin89 across the 22 Baltic Sea metagenomes with mean genome coverage greater than 30×, visualized with ‘anvi-refinè in anvi’o. Each circular layer represents one metagenome. Genome regions were ordered based on sequence composition and differential coverage using Euclidean distance and Ward linkage. The bars on the right show the total number of reads, total number of reads mapped to the genome, and percentage of reads mapped for each metagenome.

**Supplementary Figure 2.**
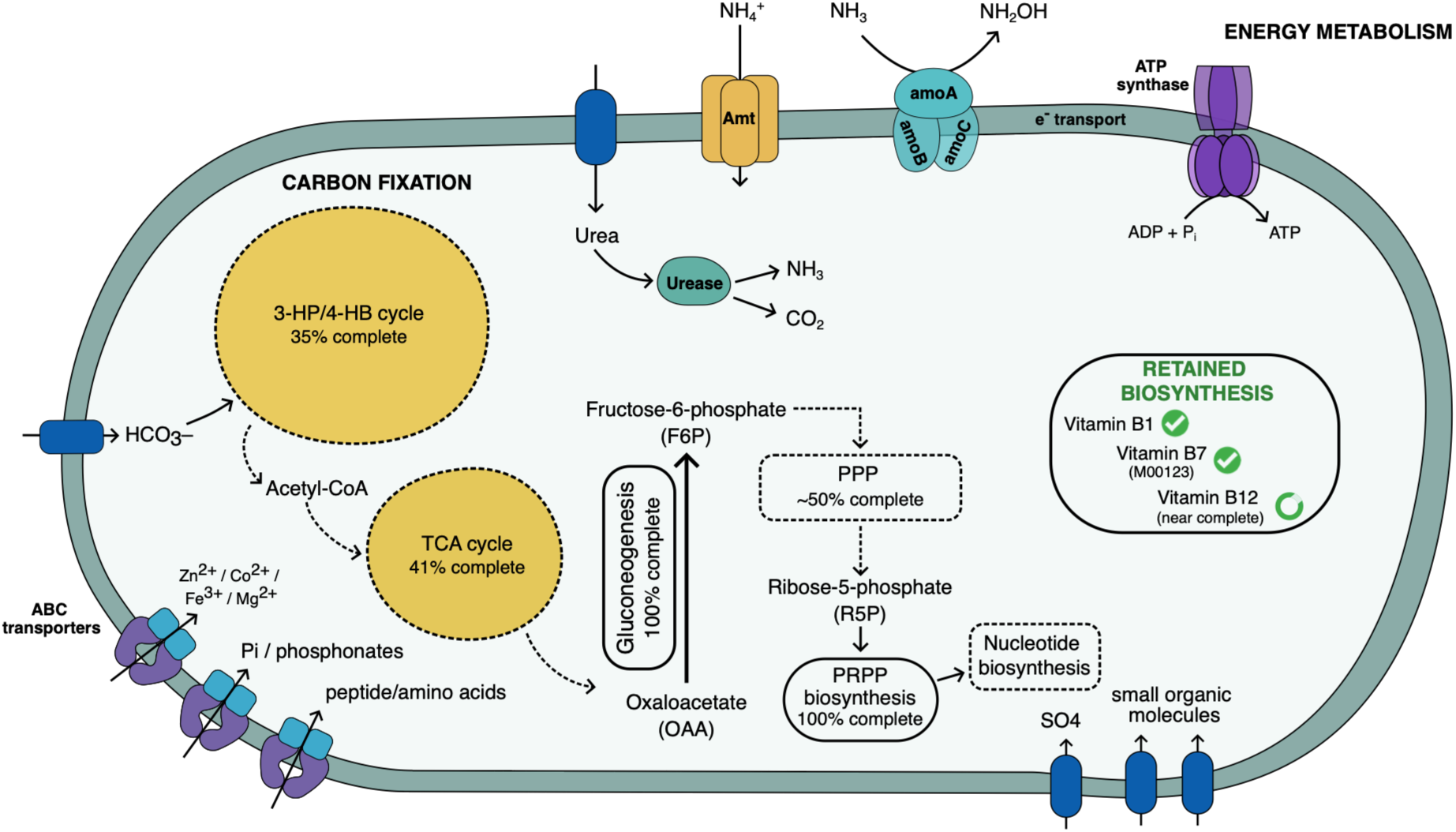
Metabolic overview of *N. supplementum* sp. nov genome. Schematic representation of energy metabolism, central carbon pathways, precursor formation, and biosynthetic capacity based on genome annotation. Functions in this archaeon include ammonia oxidation (*amoABC*), partial carbon fixation (3-HP/4-HB; 35%), and an incomplete TCA cycle reaction (41%), as well as complete gluconeogenesis and PRPP biosynthesis. Pentose phosphate pathways are partially represented (∼50%), indicating potential ribose-5-phosphate formation. B-vitamin pathways (B_1_, B_7_, and near-complete B_12_) are also shown. Transport systems support the uptake of inorganic nutrients, peptides, metals, and small organic molecules. Solid lines indicate complete pathways, and dashed lines indicate partial pathways.

## SUPPLEMENTARY NOTES

**Supplementary Note 1. Taxonomic description according to SeqCode nomenclature**

### Description of Nitrosopumilus supplementum sp. nov

*Nitrosopumilus supplementum* (sup.ple.men’tum. L. neut. n. supplementum, that with which something is made full or whole, a supply or supplement), referring to the potential of this archaeon to synthesize thiamine (B_1_), biotin (B_7_), and cobalamin (B_12_) in an environment where that combination is unusual.

The name is proposed for an archaeal species represented by a metagenome-assembled genome. The genome is classified as *Nitrosopumilus* sp001437625 according to GTDB. It shares 99.6% ANI with the previously deposited Baltic Sea genome BACL13 MAG-121220-bin23 (GCA_001437625.1). Although the two genomes were recovered from different samples, their high ANI supports their assignment to the same species-level lineage.

Metabolic annotation indicates a streamlined chemolithoautotrophic metabolism consistent with other members of *Nitrosopumilus*. The genome encodes the complete ammonia monooxygenase complex (*amoABC*), an ammonium transporter (*amt*), a putative urea transporter, and the urease complex (*ureABC*). Genes associated with carbon fixation through the 3-hydroxypropionate/4-hydroxybutyrate cycle were also detected, together with pathways supporting nucleotide biosynthesis and central carbon metabolism. The genome also retains complete biosynthetic pathways for thiamine (B_1_), all genes associated with KEGG module M00123 for biotin (B_7_), and most of the pathway for cobalamin (B_12_).

Across Baltic Sea metagenomes, the corresponding *Nitrosopumilus* population was broadly distributed and locally abundant, occurring from the northern Baltic Sea to the transition toward the North Sea and in both surface and deeper waters. Its distribution spans the strong salinity gradient characteristic of the Baltic Sea, indicating that this population is not restricted to a narrow set of Baltic environmental conditions.

The genome was recovered from the environmental sample P1994_129 (BioSample SAMEA104408620), collected on 17 June 2014 in the Western Baltic Proper at 57.1162° N, 17.6642° E, and a depth of 55.8 m. The MAG is registered in ENA as P1994_129_bin89 under BioSample SAMEA6483089 and Analysis accession ERZ1291182. The genome comprises 198 contigs, has a total size of 1404359 bp, a G+C content of 31.2%, an estimated completeness of 100%, and 0.97% contamination, all estimated with CheckM.

